# The molecular groundplan of male reproduction is partially preserved in parthenogenetic stick insects

**DOI:** 10.1101/2023.10.31.564698

**Authors:** Giobbe Forni, Barbara Mantovani, Alexander S. Mikheyev, Andrea Luchetti

## Abstract

After the loss of a trait, theory predicts that the molecular machinery underlying its phenotypic expression should decay. Yet, empirical evidence is contrasting. Here, we test the hypotheses that (1) the molecular ground plan of a lost trait could persist due to pleiotropic effects on other traits and (2) that gene co-expression network architecture could constrain individual gene expression. Our testing ground has been the *Bacillus* stick insect species complex, which contains close relatives that are either bisexual or parthenogenetic. After the identification of genes expressed in male reproductive tissues in a bisexual species, we investigated their gene co-expression network structure in two parthenogenetic species. We found that gene co-expression within the male gonads was preserved in parthenogens. Furthermore, parthenogens did not show relaxed selection on genes upregulated in male gonads in the bisexual species. As these genes were mostly expressed in female gonads, this preservation could be driven by pleiotropic interactions and an ongoing role in female reproduction. Connectivity within the network also played a key role, with highly connected - and more pleiotropic - genes within male gonad also having a gonad-biased expression in parthenogens. Our findings provide novel insight into the mechanisms which could underlie the production of rare males in parthenogenetic lineages; more generally, they provide an example of the cryptic persistence of a lost trait molecular ground plan, driven by gene pleiotropy on other traits and within their co-expression network.

**Significance:** Loss of traits commonly occurs in diverse lineages of organisms. Here we investigate what happens to genes and regulatory networks associated with these traits, using parthenogenetic insect species as a model. We investigated the fate of genes and gene regulatory networks associated with male gonads in a bisexual species in closely related parthenogens. Rather than showing signs of disuse and decay, they have been partially preserved in parthenogens. More highly pleiotropic genes in male gonads were more likely to have a gonad-biased expression profile in parthenogens. These results highlight the role of pleiotropy in the cryptic persistence of a trait molecular ground plan, despite its phenotypical absence.

## Introduction

Trait loss is a common phenomenon across the tree of life. It is predicted to have evolutionary consequences for the genes involved in the trait expression, either via drift or selection (**Hall and Colegrave 2008**). The decay of trait-specific genes after trait loss can be due to several causes, including mutations in coding sequences (**Kraaijeveld et al. 2016**) or in regulatory regions (**Sackton et al. 2019**), changes in expression patterns (**Zhang and Reed 2016**), or complete gene losses (**Suen et al. 2011**). Nonetheless, empirical evidence of decay following trait loss is often lacking. For example, parasitic wasps which have lost lipogenesis do not show sequence degradation of related genes (**Lammers et al. 2019**). Likewise, genes underlying photosynthesis are inferred to be under a strong purifying selection in a parasitic plant (**McNeal et al. 2007**). The expression of functional opsins has been observed in cave crustacean with reduced or absent eyes (**Stern and Crandall 2018**). Although these contrasting results make it difficult to generalize about the impact of trait loss on its genomic blueprint, these observations on maintenance and decay are not necessarily conflicting. Different trajectories of trait preservation or decay can be expected depending on the pleiotropic effects that its underlying genes may have on other unrelated traits. So, traits whose genes are involved in multiple biological processes are less likely to degenerate after the selective constraints on a specific trait are removed (**Smith et al. 2015**). As pleiotropy is known to affect genes expression (**Papakostas et al. 2014**; **Morandin et al. 2017**) and genes underlying complex traits have variable levels of pleiotropy (**Visscher and Yang 2016**), it is possible to hypothesize that the expression preservation of genes underlying a certain trait after its loss could be explained by their connectivity in their co-expression network topology. In fact, highly interconnected genes are more pleiotropic and more likely to be essential than others (**Jeong et al. 2001**; **MacNeil and Walhoutm, 2011**).

Parthenogenetic taxa can produce offspring without male contribution; some of these lineages only produce female individuals, thus completely lacking - or having just accidental emergence - of males (**Scali 2013**). This provides an excellent framework to unravel the consequences of trait loss. With males’ loss it can be expected that the molecular ground plan of male reproduction would degrade, and eventually be lost; yet, the rate of degradation and/or loss can be expected to depend on a plethora of phenomena, including the time since the shift in reproductive strategy, the level of pleiotropy of the underlying genes, and the rate of cryptic sex (**van der Kooi and Schwander 2014**). Our testing ground has been the *Bacillus* stick insect species complex, which includes *Bacillus grandii*, a paraphyletic group of three bisexual subspecies, and two thelytokous taxa: the obligate parthenogen *Bacillus atticus* and the facultative parthenogen *Bacillus rossius*, which includes both bisexual and parthenogenetic populations (**Scali et al. 2003**). This species complex diversified over 20 million years ago, and earlier research highlighted two cytological mechanisms of ploidy restoration in parthenogenetic taxa: in *B. atticus* the diploid egg is obtained through meiosis with central fusion by anaphase I restitution, while in *B. rossius* the embryo starts developing from an haploid egg and then diploidizes after a short while (**Scali et al. 2003**). In a previous gene expression analysis on gonadal tissues in these three taxa (**Forni et al. 2022**), genes with gonads-biased expression in parthenogens were also found to have a biased expression in male and female gonads of the bisexual species.

Here, we inferred the gene coexpression network of male gonads in the bisexual species *Bacillus grandii*; subsequently, we assessed the preservation and/or decay of expression coordination and sequence identity of the genes composing it in the two parthenogens *B. atticus* and *B. rossius*. We specifically tested the hypothesis that the expression coordination and sequence identity of genes correlated with male reproduction could be preserved due to its genes having pleiotropic effects in females (**van der Kooi and Schwander 2014**; **Schwander et al. 2013**). Furthermore, we predicted different extents of gene transcriptional preservation in parthenogens depending on their connectivity in the co-expression network, as a measure of pleiotropy.

## Results

### Characteristics of gene co-expression network in the bisexual species *B. grandii*

A weighted gene co-expression network was inferred for the bisexual species *B. grandii*, using the expression data of 2842 single-copy genes shared among the three *Bacillus* species and the outgroup *Phyllium philppinicum*. Seven co-expression modules were identified (indicated with capital letters from A to G in **Fig. 1**): three of them - D, F, and G - presented a significant, positive correlation to male gonad, and two (C and E) presented a significant and positive correlation to female gonads (**Fig. 1A**). Each module included between 105 and 909 genes, with 2.7% of genes left unassigned. Co-expression modules correlated with male and female gonads are enriched for several functions associated with transcription and meiosis, such as gene silencing, cell cycle and proliferation, morphogenesis, and chromosome organization (**Sup. Tab. 1**).

**Figure 1.**
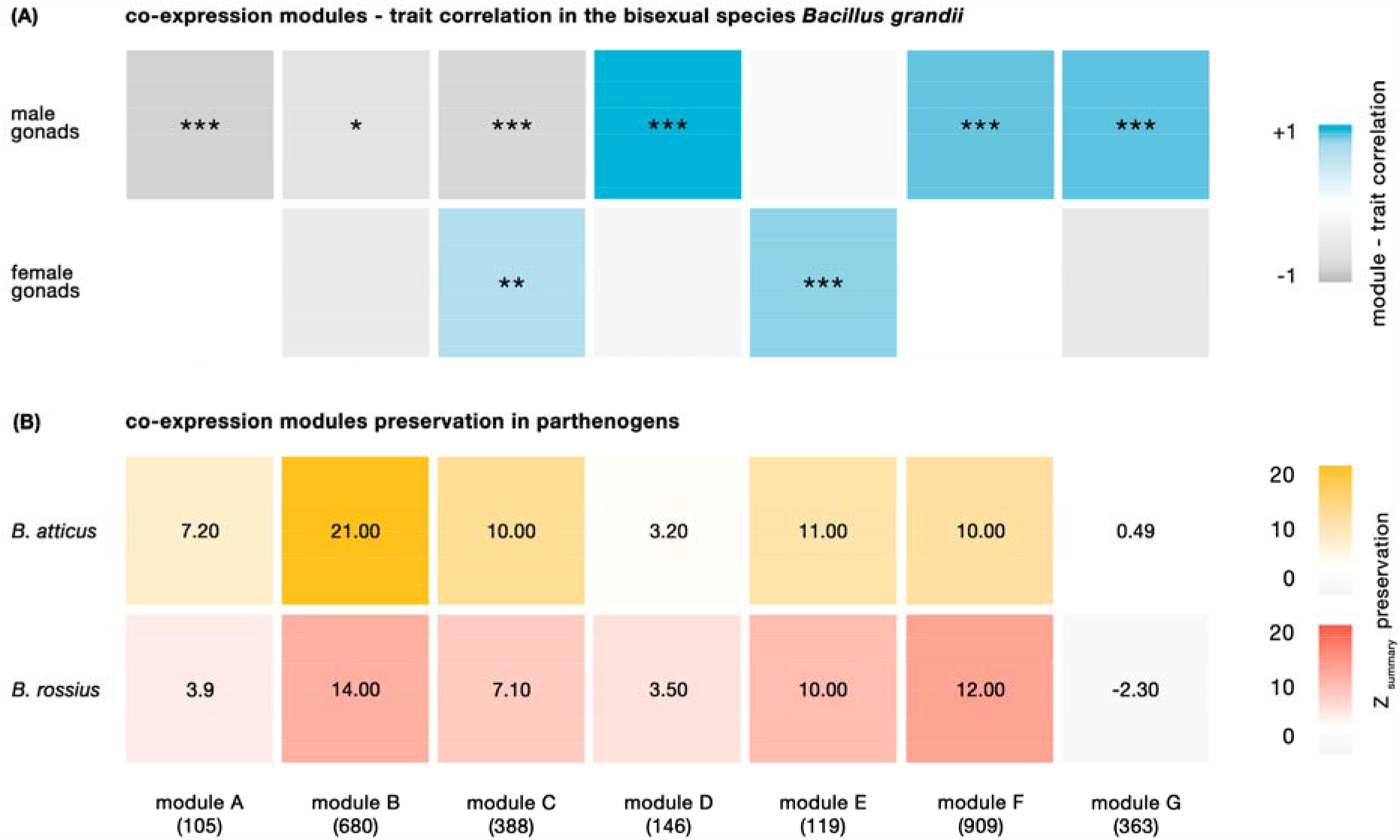
Expression preservation of male gonads-correlated genes in parthenogens. **(A)** Co-expression modules correlation to male and female gonads in the bisexual species *Bacillus grandii*. Each column corresponds to a module, while the upper and the lower rows correspond to male and female gonad trait correlation, respectively. Tiles colour intensity is proportional to module-trait correlations (*** p<0.001; ** p<0.01; * p<0.05; ns > 0.05). Module names and the number of genes composing them (in parentheses) are reported at the bottom of the figure. **(B)** Preservation of gonad-correlated modules expression coordination in the two parthenogens *Bacillus atticus* and *Bacillus rossius* gonad. Numbers in the tiles and tiles colour represtent Z_summary_ preservation: the higher the metric, the more preserved the module. Z_summary_ < 2 was considered as no evidence for module preservation, 2 < Z_summary_ < 10 as weak to moderate evidence, and Z_summary_ > 10 as strong evidence.

### Expression preservation of male gonads-correlated genes in parthenogens

After having identified gonads-correlated modules in the bisexual *B. grandii*, we tested whether network characteristics could be partially preserved in the parthenogens *B. atticus* and *B. rossius*, despite the absence of males. Following the thresholds proposed by **Langfelder et al. (2011)**, the gene network inferred in the bisexual species was found to be preserved in the two parthenogenetic species *B. atticus* and *B. rossius* (Z_summary_ preservation 15.00 for *B. rossius* and 14.00 for *B. atticus*); genes not included in the co-expression network instead didn’t show any sign of preservation (Z_summary_ preservation -0.72 for *B. rossiu*s and -0.29 for *B. atticus*). Female gonads-related modules were found consistently preserved in parthenogens, while modules correlated with male gonads exhibit different extents of preservation: module F is preserved, while module D is weakly preserved, and module G shows no signs of preservation, consistently across both parthenogens (**Fig. 1B)**.

### Sequence preservation of male gonads-correlated genes in parthenogens

Using dN/dS ratios, we tested whether male gonad-correlated genes in parthenogens experienced relaxed selection and decay in their sequence identity. No statistical difference was found in dN/dS distribution across *Bacillus* species, for all subset of genes considered (**Fig. 2A**). Within each species, both female and male gonads-correlated genes were found to have consistently undergone a weaker selective pressure with respect to gonads-unrelated genes, while there is no statistical difference between female and male gonads (**Fig. 2B**). This implies that no sign of a weakened selection is observed in male gonad related genes in parthenogens, consistent with observations on other parthenogenetic arthropods (**Brandt et al. 2017**).

**Figure 2.**
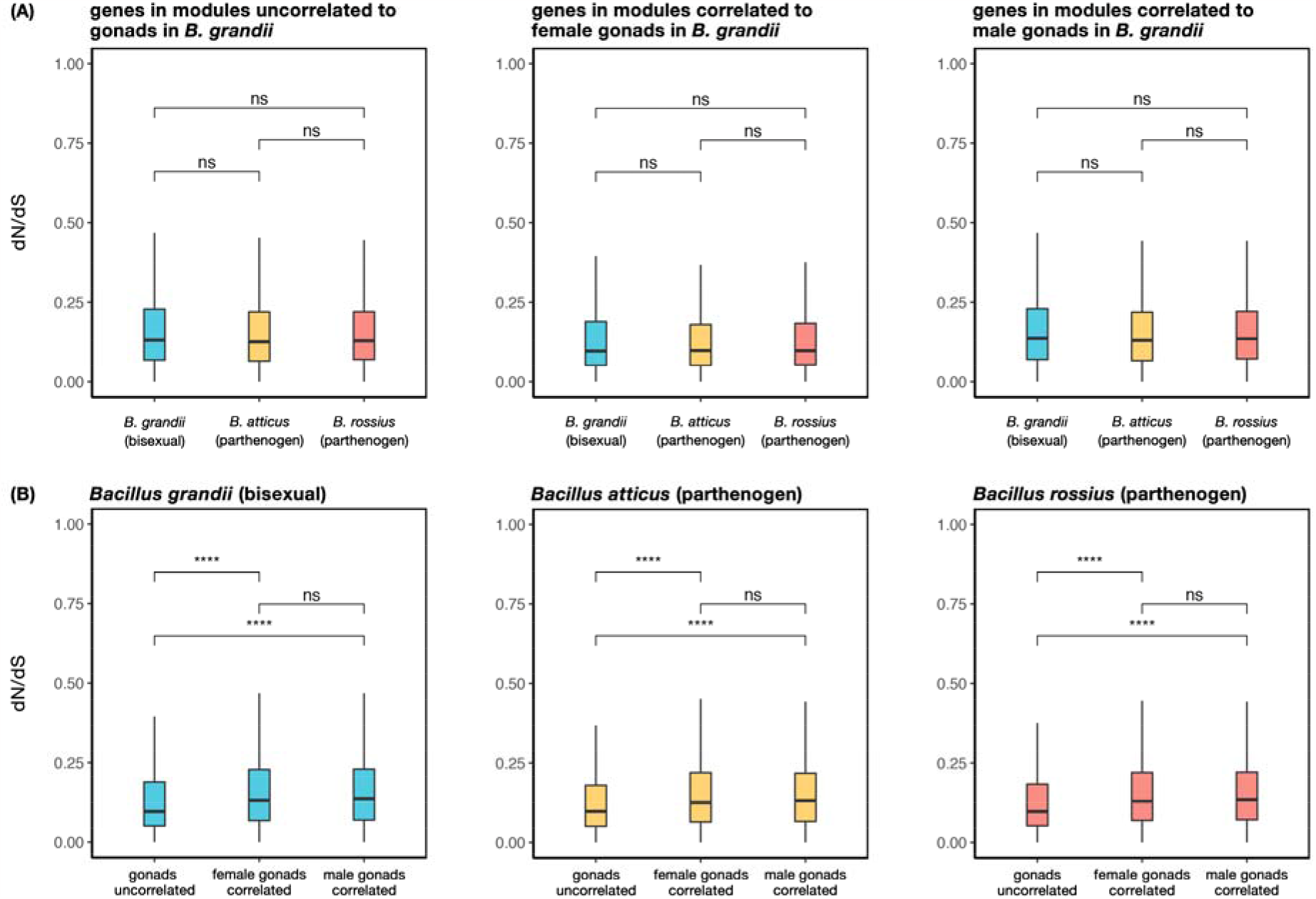
Sequence preservation of male gonad-correlated genes in parthenogens. **(A)** Cross-species comparison of dN/dS ratios for gonads-uncorrelated, female gonads-correlated and male gonads-correlated genes. **(B)** Within-species comparison of dN/dS ratios for gonads-uncorrelated, female gonads-correlated and male gonads-correlated genes (*** p<0.001; ** p<0.01; * p<0.05; ns > 0.05). Overall, no signature of relaxed selection was found in the sequence evolution of male gonad-correlated genes in parthenogens.

### Pleiotropy of male gonads-correlated genes on other traits and within the co-expression network

We then focused on the interplay between gene gonads-biased pattern of expression in parthenogens and pleiotropic interactions on other traits and within their co-expression network. In the bisexual species *B. grandii*, genes found in co-expression modules positively correlated with female gonads are found to have a gonads-biased expression in female gonads, including parthenogenetic ones, but they appear strongly downregulated in male ones (**Fig. 3A**). Conversely, most of the genes found in co-expression modules positively correlated with male gonads have a gonads-biased expression in both bisexual males and females, and parthenogenetic females (**Fig. 3B**). Thus, the preservation of connectivity and density of most of the male gonad genes co-expression network in parthenogens may be explained by pleiotropic effects in both sexes of genes underlying male gonads physiological pathways, as previously hypothesized (**van der Kooi and Schwander 2014**; **Schwander et al. 2013**). Interestingly, a large part of male gonads-correlated genes has a gonads-biased profile of expression in parthenogenetic females, and their connectivity in *B. grandii* co-expression network was found to have a significant and positive correlation with parthenogens LogFC (*B. atticus*: Spearman correlation: r=0.34, p<0.001; *B. rossius*: r=0.31, p<0.001; **Fig. 3B**). This suggests that highly connected genes correlated with male gonads are most likely to show a gonads-biased transcription pattern in parthenogens, while loosely connected ones lack a gonad-specific expression in parthenogens.

**Figure 3.**
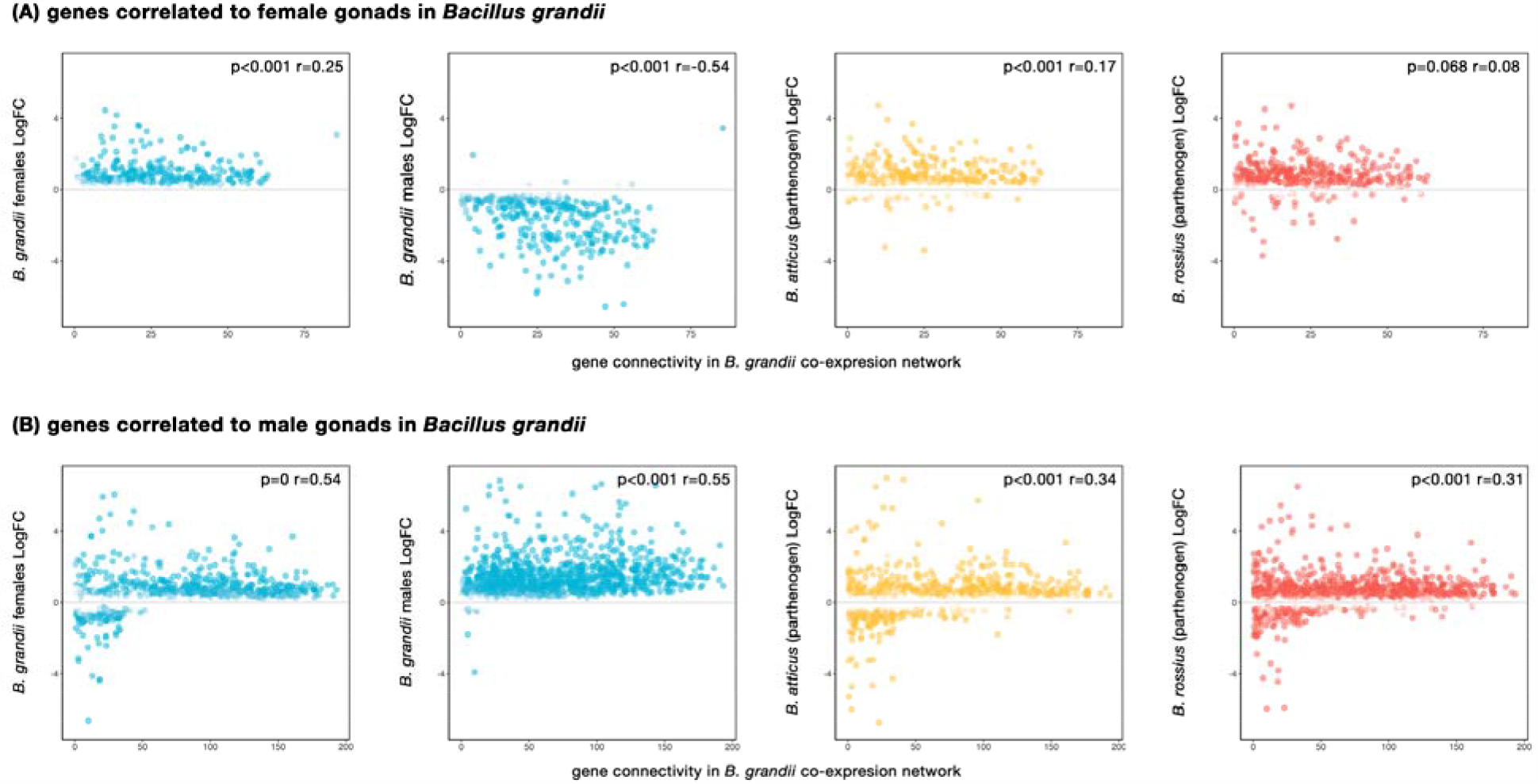
Pleiotropy of male gonads-correlated genes on other traits and within the co-expression network. **(A)** Spearman correlation between gene connectivity in the bisexual *Bacillus grandii* co-expression network and the LogFC between reproductive and non-reproductive tissues in each *Bacillus* species and sex, for genes which were found to be correlated to female gonads in the bisexual *Bacillus grandii*; female gonad genes are found to be generally downregulated in male gonad. **(B)** Spearman correlations between gene connectivity in the bisexual *Bacillus grandii* co-expression network and the LogFC between reproductive and non-reproductive tissues in each *Bacillus* species and sex, for genes which were found to be correlated to male gonads in the bisexual *Bacillus grandii*. male gonad genes are found to be generally upregulated in female and parthenogens’ gonads; furthermore, genes with higher connectivity are more likely to be expressed in female and parthenogens’ gonads. Points colour is based on differential expression FDR.

## Discussion

We examined the fate of the male gonads gene regulatory network in parthenogenetic taxa. Our results reveal the persistence of its ground plan in both transcription coordination and sequence identity. These findings are coherent with trait loss being coupled mainly with transcriptional modifications (**Roscito et al. 2018**; **Smith et al. 2015**) and changes affecting a few genes (**Zhang and Reed 2016**; **Yagound et al. 2020**), thus preserving a large portion of the trait blueprint after its phenotypic loss (**Smith et al. 2015**; **Stern and Crandall 2018**; **Bast et al. 2018**; **Lammers et al. 2019**). As for other lost traits, the persistence of male reproduction ground plan in *Bacillus* parthenogens can have two non-mutually exclusive explanations: (1) neutral decay necessitating extended time periods to effect measurable loss of function and/or (2) physiological pathways consisting of pleiotropic genes, which are under selection for other traits or for their essentiality.

Our results support a role for gene pleiotropy on other traits and within the co-expression network in the observed preservation. Highly connected genes are more likely to have pleiotropic effects also for functions that are not related to the lost trait, with respect to loosely connected ones **(Tyler et al. 2009**; **Wang et al. 2010**), and pleiotropy is also known to affect genes expression (**Papakostas et al. 2014**; **Morandin et al. 2017**); theoretical and empirical observations also predict a strong correlation between connectivity and essentiality, so that the degradation of genes largely responsible for maintaining network connectivity could be detrimental or even lethal (**Jeong et al. 2001**). Furthermore, our results provide a possible explanation for cryptic sex and rare males (i.e., episodic generation of males and bisexual reproduction). There is an increasing number of instances where signatures of cryptic sex are found in “ancient” parthenogens (**Vakhrusheva et al. 2020**; **Boyer et al. 2021**; **Freitas et al. 2023**); at the same time, parthenogenetically produced males have often been observed in many species, and have also been reported for *B. atticus* (**Scali 2013**). Our findings are also compatible with the observed masculinization of sex-biased gene expression following the shift to parthenogenesis (**Parker et al. 2019**) but highlight, instead, the role of prior constraints (gene pleiotropy and essentiality) alongside selective forces (**Losos et al. 2011**).

However, while *Bacillus* species diverged several million years ago (**Scali et al. 2003**), it is impossible to date when the shift in reproductive strategy happened and whether there has been enough for the neutral decay to be observed. However, a simple model of neutral decay would not predict some of our findings, such as greater involvement of pleiotropic male gonads genes in the ovaries of asexual taxa. Thus, we believe this hypothesis is a less likely explanation for our data, though it cannot be strictly ruled for some aspects of our findings, such as the lack of relaxed selection on male gonads-upregulated genes in the asexual species.

A large body of literature focuses on how traits are established (**Fisher et al. 2020**; **Almudi et al. 2020**); nonetheless unraveling the processes associated with trait cryptic persistence can also provide a broader insight into the selective forces that underlie trait evolution (**Bruce and Patel 2022**). The reversal to an ancestral state after trait loss is considered an unlikely event (e.g., Dollo’s law), as it is expected to be coupled with the decay of the trait molecular ground plan (**Marshall et al. 1994**). Yet, numerous examples of such reversals have been proposed, including oviparity in lizards (**Esquerré et al. 2020**), insects wings (**Forni et al. 2022**), and sex itself (**Domes et al. 2007**); the results presented here suggest a possible mechanism which could underlie the reversion to a once-lost ancestral state.

## Materials and Methods

De novo reference transcriptome assemblies and annotation, differential expression analyses between reproductive and non-reproductive tissues, and orthology inference were generated previously (**Forni et al. 2022**). A weighted gene co-expression network was inferred for the bisexual species *B. grandii*. This approach identifies genes with coordinated expression patterns across samples, using a hierarchical clustering approach; genes are clustered into modules of highly interconnected genes which are then correlated with specific traits (**Stuart et al. 2003**). Using the gene level abundance estimates of 24 *B. grandii* samples (6 replicates of legs and gonads tissues, for male and female specimens), a matrix of TPM expression values was constructed and cross-sample normalized using the TMM method implemented in edgeR v3.40.0 (**Robinson and Oshlack 2010**). The analysis was performed using the R package WGCNA v1.71 (**Langfelder and Horvath 2008**). A signed network was inferred based on the criterion of approximate scale-free topology, using the blockwiseModules WGCNA function, a soft thresholding power of 19, and a dendrogram cut height of 0.2, modules defined to contain at least 30 genes. All genes found in modules that had a positive significance correlation with male gonad are subsequently defined as male gonads-correlated, while all genes inside modules that had a positive significance with female gonad are defined as female gonads-correlated; all genes unrelated to male or female gonads were considered separately. GO-terms were generated for each species using Argot 2.5 with a TotalScore > 200 (**Lavezzo et al. 2016**) and subsequently associated with each OG, collapsing multiple entries of the same term. Each module enrichment analysis were performed with the TopGO v2.32.0 package in Bioconductor, using Fisher exact test and both elim and weight algorithms - which consider GO hierarchy (**Alexa and Rahnenführer 2009**). GO-terms were considered significantly enriched when elim p<□0.05.

Leveraging 6 gonads and 6 legs samples for each species, module preservation statistics were inferred in WGCNA using the modulePreservation function, with 500 permutations and assuming a signed network (**Langfelder et al. 2011**). The Z_summary_ metric has been used to quantify the preservation of density and connectivity patterns of the bisexual species modules in the species which shifted to parthenogenesis.

For each orthogroup, protein sequences were aligned using MAFFT v7.520 (**Katoh and Standley 2013**) with default parameters and retro-translated to a nucleotide alignment using pal2nal v14.1 (**Suyama et al. 2006**). For each alignment, the species tree branch lengths were optimized in a maximum-likelihood framework, using a codon-aware GTR+G model, with RaxML v8.2.12 (**Stamatakis 2014**). dN/dS ratios were calculated along the branches of the species phylogeny using codeml v4.10.6 (**Yang 2007**): a model with one dN/dS for all branches of the tree and a model with a branch-specific dN/dS were compared using a Likelihood Ratio Test (LRT), with p-values adjusted for multiple comparisons using Benjamini-Hochberg’s correction in R. dN/dS values for each terminal branch leading to the three *Bacillus* taxa were retrieved for each orthogroup; dN/dS values associated to dN>3 (implying substitutions saturation) and artefactual dN/dS values of 999 were excluded. Furthermore, dN/dS > 1, were excluded in order to assess the strength of purifying selection without considering confounding instances of positive selection. Statistical comparisons of different dN/dS rations were carried out using the Wilcoxon non-parametric test.

Using the same parameters used for network construction, gene connectivity has been inferred for each gene in the bisexual species gene co-expression network with the intramodularConnectivity WGCNA function; this metric is intended as a measure of how correlated the expression of a gene is with others and as a proxy for the extent of a gene’s pleiotropic interactions. Using R (4.0.2; **R Core Team, 2017**), Spearman correlations were calculated between gene total connectivity (kTotal) in the bisexual *B. grandii* co-expression network and the LogFC found between reproductive and non-reproductive tissues of the two *Bacillus* parthenogens, male and female gonad-correlated genes.

## Data availability

All RNA-seq reads are deposited on the NCBI SRA database under acc. nos. SRX7034623–SRX7034670 (Bioproject: PRJNA578804). Assembled transcriptomes have been uploaded to the NCBI TSA database under acc. nos GJDY01000000, GJDZ01000000, and GJEA00000000. All custom scripts and intermediate files are available at https://github.com/for-giobbe/bacillus_cryptic_persistence.

## Funding

This work was supported by the Canziani funding to AL and BM.

## Acknowledgments

We thank Giovanni Piccinini and Fabrizio Ghiselli for the valuable comments on the draft. We thank Alex Cussigh for helping to rear *Bacillus* insects.

## Notes

### Competing Interest Statement

The authors have declared no competing interest.

## References

1. Alexa A, Rahnenfuhrer J (2019). topGO: Enrichment Analysis for Gene Ontology. R package version 2.38.1.

2. Almudi I, Vizueta J, Wyatt CD, de Mendoza A, Marlétaz F, Firbas PN, Feuda R, Masiero G, Medina P, Alcaina-Caro A, Cruz F. 2020. Genomic adaptations to aquatic and aerial life in mayflies and the origin of insect wings. Nature communications, 11(1), pp.1–11.

3. Bast J, Parker DJ, Dumas Z, Jalvingh KM, Tran Van P, Jaron KS, Figuet E, Brandt A, Galtier N, Schwander T. 2018. Consequences of asexuality in natural populations: insights from stick insects. Molecular biology and evolution, 35(7), pp.1668–1677.

4. Boyer L, Jabbour-Zahab R, Mosna M, Haag CR, Lenormand T. 2021. Not so clonal asexuals: Unraveling the secret sex life of Artemia parthenogenetica. Evolution Letters, 5:164–174.

5. Brandt A, Schaefer I, Glanz J, Schwander T, Maraun M, Scheu S, Bast J. 2017. Effective purifying selection in ancient asexual oribatid mites. Nature communications, 8(1), pp.1–9.

6. Bruce HS, Patel NH. 2022. The Daphnia carapace and other novel structures evolved via the cryptic persistence of serial homologs. Current Biology, 32(17), pp.3792–3799.

7. Domes K, Norton RA, Maraun M, Scheu S. 2007. Reevolution of sexuality breaks Dollo’s law. Proceedings of the National Academy of Sciences, 104(17), pp.7139–7144.

8. Fisher CR, Wegrzyn JL, Jockusch EL. 2020. Co-option of wing-patterning genes underlies the evolution of the treehopper helmet. Nature Ecology & Evolution, 4(2), pp.250–260.

9. Forni G, Martelossi J, Valero P, Hennemann FH, Conle O, Luchetti A, Mantovani B. 2022. Macroevolutionary analyses provide new evidence of phasmid wings evolution as a reversible process. Systematic Biology, 71(6), pp.1471–1486.

10. Forni G, Mikheyev AS, Luchetti A, Mantovani B. 2022. Gene transcriptional profiles in gonads of Bacillus taxa (Phasmida) with different cytological mechanisms of automictic parthenogenesis. Zoological Letters, 8(1), pp.1–14.

11. Freitas S, Parker DJ, Labédan M, Dumas Z, Schwander T. 2023. Evidence for cryptic gene flow in parthenogenetic stick insects of the genus Timema. BioRxiv, pp.2023–01.

12. Esquerré D, Brennan IG, Catullo RA, Torres-Pérez F,, Keogh JS. 2019. How mountains shape biodiversity: The role of the Andes in biogeography, diversification, and reproductive biology in South America’s most species-rich lizard radiation (Squamata: Liolaemidae). Evolution, 73(2), pp.214–230.

13. Hall AR, Colegrave N. 2008. Decay of unused characters by selection and drift. Journal of Evolutionary Biology, 21(2), pp.610–617.

14. Jeong H, Mason SP, Barabási AL, Oltvai ZN. 2001. Lethality and centrality in protein networks. Nature, 411, 41–42.

15. Katoh K, Standley DM. 2013. MAFFT multiple sequence alignment software version 7: improvements in performance and usability. Molecular biology and evolution, 30(4), pp.772–780.

16. Kraaijeveld K, Anvar SY, Frank J, Schmitz A, Bast J, Wilbrandt J, Petersen M, Ziesmann T, Niehuis O, De Knijff P, Den Dunnen JT. 2016. Decay of sexual trait genes in an asexual parasitoid wasp. Genome biology and evolution, 8(12), pp.3685–3695.

17. Langfelder P, Horvath S. 2008. WGCNA: an R package for weighted correlation network analysis. BMC bioinformatics, 9(1), p.559.

18. Langfelder P, Luo R, Oldham MC, Horvath S. 2011. Is my network module preserved and reproducible?. PLoS Computational Biology, 7(1), p.e1001057.

19. Lammers M, Kraaijeveld K, Mariën J, Ellers J. 2019. Gene expression changes associated with the evolutionary loss of a metabolic trait: lack of lipogenesis in parasitoids. BMC genomics, 20(1), pp.1–14.

20. Lavezzo E, Falda M, Fontana P, Bianco L, Toppo S. 2016. Enhancing protein function prediction with taxonomic constraints–The Argot2. 5 web server. Methods, 93, pp.15–23.

21. Losos JB. 2011. Convergence, adaptation, and constraint. Evolution, 65(7), pp.1827–1840.

22. MacNeil LT, Walhout AJ. 2011. Gene regulatory networks and the role of robustness and stochasticity in the control of gene expression. Genome research, 21(5), pp.645–657.

23. Marshall CR, Raff EC, Raff RA. 1994. Dollo’s law and the death and resurrection of genes. Proceedings of the National Academy of Sciences, 91(25), pp.12283–12287.

24. Morandin C, Mikheyev AS, Pedersen JS, Helanterä H. 2017. Evolutionary constraints shape caste-specific gene expression across 15 ant species. Evolution, 71(5), pp.1273–1284.

25. McNeal JR, Kuehl JV, Boore JL, de Pamphilis CW. 2007. Complete plastid genome sequences suggest strong selection for retention of photosynthetic genes in the parasitic plant genus Cuscuta. BMC Plant Biology, 7(1), p.57.

26. Papakostas S, Vøllestad LA, Bruneaux M, Aykanat T, Vanoverbeke J, Ning M, Primmer CR, Leder EH. 2014. Gene pleiotropy constrains gene expression changes in fish adapted to different thermal conditions. Nature Communications, 5(1), pp.1–9.

27. Parker DJ, Bast J, Jalvingh K, Dumas Z, Robinson-Rechavi M, Schwander T. 2019. Sex-biased gene expression is repeatedly masculinized in asexual females. Nature communications, 10(1), pp.1–11.

28. R. Core Team. (R: A Language and Environment for Statistical Computing, 2017).

29. Robinson MD, Oshlack A. 2010. A scaling normalization method for differential expression analysis of RNA-seq data. Genome biology, 11(3), pp.1–9.

30. Roscito JG, Sameith K, Parra G, Langer BE, Petzold A, Moebius C, Bickle M, Rodrigues MT, Hiller M. 2018. Phenotype loss is associated with widespread divergence of the gene regulatory landscape in evolution. Nature communications, 9(1), pp.1–15.

31. Sackton TB, Grayson P, Cloutier A, Hu Z, Liu JS, Wheeler NE, Gardner PP, Clarke JA, Baker AJ, Clamp M, Edwards SV. 2019. Convergent regulatory evolution and loss of flight in paleognathous birds. Science, 364(6435), pp.74–78.

32. Scali V, Passamonti M, Marescalchi O, Mantovani B, 2003. Linkage between sexual and asexual lineages: genome evolution in Bacillus stick insects. Biological Journal of the Linnean Society. 2003;79:137–50.

33. Scali V. 2013. A “ghost” phasmid appearance: the male Bacillus atticus (Insecta: Phasmatodea). Italian Journal of Zoology, 80(2), pp.227–232.

34. Schwander T, Crespi BJ, Gries R, Gries G. 2013. Neutral and selection-driven decay of sexual traits in asexual stick insects. Proceedings of the Royal Society B: Biological Sciences, 280(1764), p.20130823.

35. Smith CR, Helms Cahan, S., Kemena, C., Brady, S.G., Yang, W., Bornberg-Bauer, E., Eriksson, T., Gadau, J., Helmkampf, M., Gotzek, D., Okamoto Miyakawa, M., Suarez, A.V ., Mikheyev A., 2015. How do genomes create novel phenotypes? Insights from the loss of the worker caste in ant social parasites. Molecular biology and evolution, 32(11), pp.2919–2931.

36. Stamatakis A. 2014. RAxML version 8: a tool for phylogenetic analysis and post-analysis of large phylogenies. Bioinformatics, 30(9), pp.1312–1313.

37. Stern DB, Crandall KA. 2018. The evolution of gene expression underlying vision loss in cave animals. Molecular biology and evolution, 35(8), pp.2005–2014.

38. Stuart JM, Segal E, Koller D, Kim SK. 2003. A gene-coexpression network for global discovery of conserved genetic modules. Science, 302(5643), pp.249–255.

39. Suen G, Teiling C, Li L, Holt C, Abouheif E, Bornberg-Bauer E, Bouffard P, Caldera EJ, Cash E, Cavanaugh A, Denas O. 2011. The genome sequence of the leaf-cutter ant Atta cephalotes reveals insights into its obligate symbiotic lifestyle. PLoS Genet, 7(2), p.e1002007.

40. Suyama M, Torrents D, Bork P. 2006. PAL2NAL: robust conversion of protein sequence alignments into the corresponding codon alignments. Nucleic acids research, 34(suppl_2), pp.W609–W612.

41. Tyler AL, Asselbergs FW, Williams SM, Moore JH. 2009. Shadows of complexity: what biological networks reveal about epistasis and pleiotropy. Bioessays, 31(2), pp.220–227.

42. van der Kooi CJ, Schwander T. 2014. On the fate of sexual traits under asexuality. Biological Reviews, 89(4), pp.805–819.

43. Vakhrusheva OA, Mnatsakanova EA, Galimov YR, Neretina TV, Gerasimov ES, Naumenko SA, Ozerova SG, Zalevsky AO, Yushenova IA, Rodriguez F, Arkhipova IR, Penin AA, Logacheva MD, Bazykin GA, Kondrashov AS. 2020. Genomic signatures of recombination in a natural population of the bdelloid rotifer Adineta vaga. Nature Communication 11:6421.

44. Visscher PM, Yang J. 2016. A plethora of pleiotropy across complex traits. Nature genetics, 48(7), pp.707–708.

45. Wang Z, Liao BY, Zhang J. 2010. Genomic patterns of pleiotropy and the evolution of complexity. Proceedings of the National Academy of Sciences, 107(42), pp.18034–18039.

46. Yagound B, Dogantzis KA, Zayed A, Lim J, Broekhuyse P, Remnant EJ, Beekman M, Allsopp MH, Aamidor SE, Dim, O, Buchmann G. 2020. A Single Gene Causes Thelytokous Parthenogenesis, the Defining Feature of the Cape Honeybee Apis mellifera capensis. Current Biology.

47. Yang Z. 2007. PAML 4: phylogenetic analysis by maximum likelihood. Molecular biology and evolution, 24(8), pp.1586–1591.

48. Zhang L, Reed RD. 2016. Genome editing in butterflies reveals that spalt promotes and Distal-less represses eyespot colour patterns. Nature communications, 7(1), pp.1–7.

